# High-throughput Kinetics using Capillary Electrophoresis and Robotics (HiKER) platform used to Study T7, T3, and Sp6 RNA Polymerase Misincorporation

**DOI:** 10.1101/2024.03.20.585964

**Authors:** Zachariah I. Carter, William B. O’Brien, Sean Lund, Andrew F. Gardner

## Abstract

T7 RNA Polymerase (RNAP) is a well-studied and widely used enzyme with recent applications in the production of RNA vaccines. For over 50 years denaturing sequencing gels have been used as a key analysis tool for probing the kinetic mechanism of T7 RNAP nucleotide addition. However, sequencing gels are both slow and low throughput limiting their utility for comprehensive enzyme analysis. Here, we report the development of HiKER; (<u>Hi</u>gh-throughput <u>K</u>inetics using Capillary <u>E</u>lectrophoresis and <u>R</u>obotics) a high-throughput pipeline to quantitatively measure enzyme kinetics. We adapted a traditional polymerase misincorporation assay for fluorescent detection at scale allowing rapid estimates of RNAP misincorporation in different experimental conditions. In addition, high-throughput kinetics reactions were automated using an open-source OT-2 liquid handling robot. The platform allows multiple weeks’ worth of data to be collected in mere days. Using this platform, ∼1500 time points were collected in a single workday. T7 RNAP exhibited dramatic differences in both observed rate constant and amplitude depending on the mismatch examined. An average misincorporation frequency of ∼45 misincorporations per million bases was estimated using HiKER and is consistent with previous observations from next generation sequencing studies. Misincorporation time courses for T3 RNAP and Sp6 RNAP were similar to T7 RNAP suggesting conserved kinetic mechanisms. Interestingly, dramatic changes in the extent of misincorporation were observed in the three RNAPs depending on the mismatch. Extension from base mismatch experiments showed differences between T7, T3, and Sp6 RNAP. Sp6 RNAP was the slowest to extend from a mismatch followed by T7 RNAP and then T3 RNAP. Taken together the results presented here demonstrate the capabilities of HiKER to carry out high-throughput enzymology studies. Importantly, this pipeline and the corresponding analysis strategies are affordable, open-source, and broadly applicable to many enzymes.

## Introduction

The central dogma of molecular biology states genetic information flows in one direction, with DNA transcribed to RNA followed by translation to protein. The transfer of information is extremely accurate in all organisms as mistakes are often detrimental [1-4]. DNA-dependent RNA synthesis by RNA polymerase (RNAP) is an exceptionally precise process with a single misincorporation event occurring after ∼100,000 correct nucleotide additions (50 misincorporations per million bases) [5-7]. T7 RNA polymerase (RNAP) is a single subunit RNA polymerase whose native activity is involved in the T7 phage infection process of *Escherichia coli*. This enzyme has been extensively studied for the past 50 years [8-14] and continues to be the subject of active research both for biotechnology applications and as a model RNA polymerase [15-22]. T7 RNAP was the major RNAP utilized to synthesize the mRNA transcripts found in billions of doses of the novel mRNA COVID-19 mRNA vaccine [23, 24].

Transcription by RNAPs can be divided into three distinct steps: initiation, elongation, and termination. Initiation is the process in which the polymerase binds with extreme specificity to a promoter sequence and incorporates a short RNA transcript to enable elongation [25-28]. Following initiation, elongation proceeds and enables numerous rounds of nucleotide addition to the nascent RNA transcript. Termination is the last step and is the process by which the polymerase halts and dissociates [29-31]. For scientific studies, promoter independent transcription experiments are often performed using a minimal scaffold (or transcription bubble) to mimic processive elongation [32-41].

Synthesis of correct transcripts is critical for all RNAPs. Phage synthesis of the correct RNA sequence is critical to the life cycle of the virus. Remarkably, phage RNAPs can synthesize∼20,000 nucleotide long RNA molecules without making a single error [8, 42]. Unlike other multi-subunit RNAPs found in both prokaryotes and eukaryotes, phage T7 RNAP has no known proofreading activity [43-47]. Consequently, synthesis of the correct transcript requires T7 RNAP to exhibit extremely high fidelity that relies on low levels of misincorporation.

Kinetic assays monitoring DNA/RNA length have been widely used to understand polymerase nucleotide addition [7, 38, 40, 41, 48-52]. The overwhelming majority of *in vitro* kinetic studies measuring polymerase nucleotide addition/misincorporation utilize polyacrylamide gel electrophoresis (PAGE) [7, 32, 34, 37-41, 49, 53-55]. PAGE has many practical benefits including simplicity, affordability and requiring minimal experimental equipment. Unfortunately this methodology is also tedious and low throughput, with a single PAGE system only processing a maximum of ∼50 time points a day. Thus, comprehensive comparisons across enzymes and experimental conditions is hindered due to these experimental limitations.

A complementary method for quantifying misincorporation is the counting of misincorporations in fully synthesized transcripts using next generational sequencing (NGS) [5, 6, 56]. NGS approaches are incredibly useful as they allow probing of many different sequence contexts. Reported values using <u>S</u>ingle-<u>M</u>olecule <u>R</u>eal-<u>T</u>ime (SMRT) sequencing combined T7 and reverse transcriptase misincorporation at ∼50 errors per million bases [5, 6]. It should be highlighted that SMRT sequencing relies on a reverse transcription step and the reported misincorporation is the sum of both T7 RNAP and reverse transcriptase misincorporation. A general drawback to all NGS methods is the loss of mechanistic information as only the final RNA product is observed. Additionally, NGS methods require expensive specialized reagents and equipment. Consequently, alternative methods are necessary when characterizing kinetic or mechanistic differences between experimental conditions.

Previous PAGE kinetic studies of T7 RNAP misincorporation have shown pyrimidine-purine mismatches to be more common compared to pyrimidine-pyrimidine or purine-purine [7, 34, 35]. Time courses were fit to a single exponential resulting in an observed rate constant for misincorporation and an amplitude. The reported observed misincorporation rate constants are slow (at less than ∼0.2 min^-1^) with amplitudes of ∼0.75 [7, 34, 35, 57]. Correct nucleotide addition, in contrast, is several orders of magnitude faster with values ∼12,000 min^-1^ or ∼200 s^-1^ (**Fig.1**) [33, 57]. Consequently, correct nucleotide addition is kinetically favorable compared to misincorporation. The ratio of these two rate constants suggests a misincorporation frequency of∼20 misincorporations per million nucleotide incorporations and is in agreement with measurements made using NGS[5, 6].

T3 and Sp6 are two other single subunit phage RNAPs. Fewer publications exist for T3 and Sp6 RNAP when compared to the breath of studies of T7 RNAP. A high degree of structural similarity has been observed for T7, T3, and Sp6 RNAP [58, 59]. Similar to T7 RNAP, T3 RNAP has shown a single nucleotide addition observed rate constant of ∼170 s^-1^ [60]. However, the nucleotide misincorporation kinetics for T3 and Sp6 RNAP are currently unexplored.

Previously we demonstrated how many biochemical characterizations can be analyzed using capillary electrophoresis [61]. In this study we expanded that approach by developing a platform merging robotics with capillary electrophoresis. The high-throughput kinetics platform HiKER (<u>Hi</u>gh-throughput <u>K</u>inetics using Capillary <u>E</u>lectrophoresis and Robotics) was developed and utilized to examine the misincorporation properties of T7, T3, and Sp6 RNAP (**Fig. 2**). A promoter independent transcription assay was used to efficiently monitor individual nucleotide misincorporations. Additionally, HiKER was used to probe the kinetics of T7, T3, and Sp6 RNAP extension from a base mismatch. HiKER dramatically increased the feasibility of examining the misincorporation properties of multiple RNAPs compared to traditional sequencing gels. A comparison of T7, T3, and Sp6 RNAP revealed conserved misincorporation profiles between the enzymes suggesting similar mechanisms of nucleotide misincorporation. The platform is broadly applicable and easily adaptable for many enzymology studies.

**Fig. 1.**
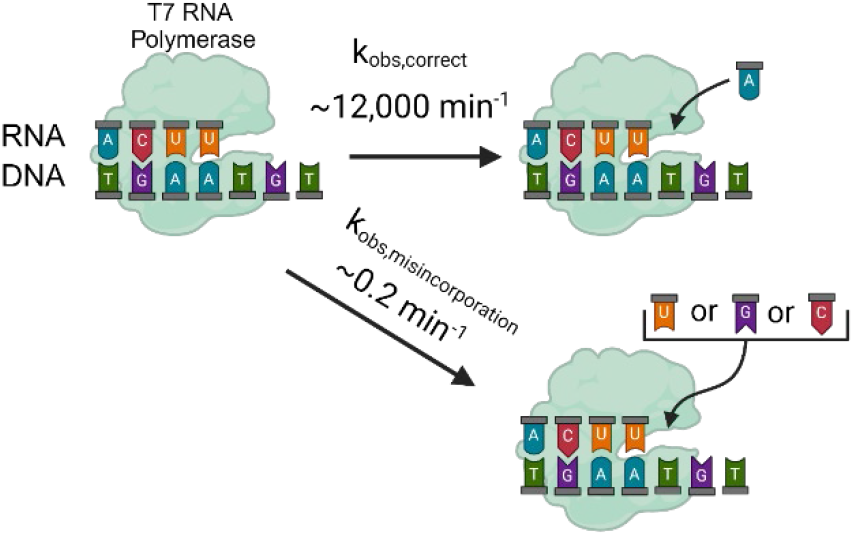
Illustration of branching ratio between T7 RNAP correct nucleotide addition and incorrect nucleotide addition. Image was generated using BioRender.

**Fig. 2.**
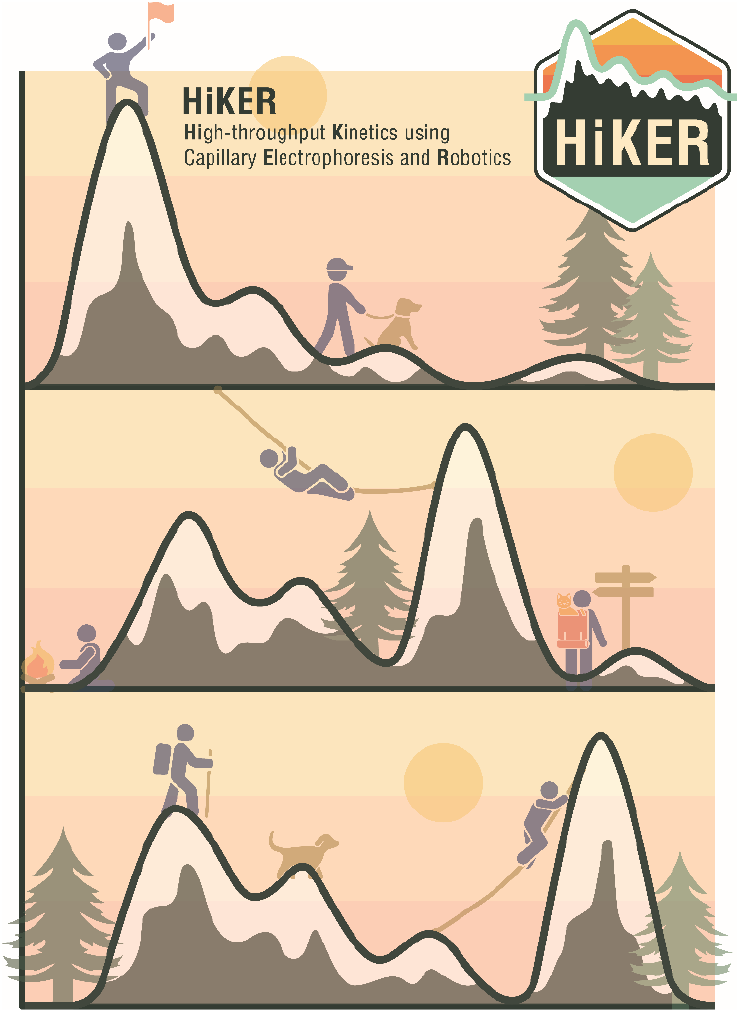
HiKER logo and illustration. The individual mountains are symbolic of the capillary electrophoresis traces. Three are shown to illustrate the throughput capabilities of HiKER. The movement from left to right of the largest peak is meant to convey the progression of a chemical reaction moving from substrate to larger products.

## Results

### In vitro RNA Polymerase Misincorporation Assay

For this study, T7 RNAP misincorporation was chosen as a proof-of-principle model system to use with HiKER as T7 RNAP has been well studied [7, 35, 57]. McAllister’s group previously demonstrated that T7 RNAP will extend from a variety of promoter independent primer/template scaffold designs [32]. In contrast to a cellular elongation complex, a minimal scaffold was selected for this study consisting of a template DNA strand and a 5’ fluorophore (FAM) labeled RNA primer (**Fig. 3A**). Importantly, the minimal scaffold shown here has been shown to exhibit properties of promoter-initiated elongation complexes [32]. The simplicity of this elongation complex substrate complemented the high-throughput properties of HiKER. Elongation complexes were rapidly assembled and the misincorporation properties were investigated.

**Fig. 3.**
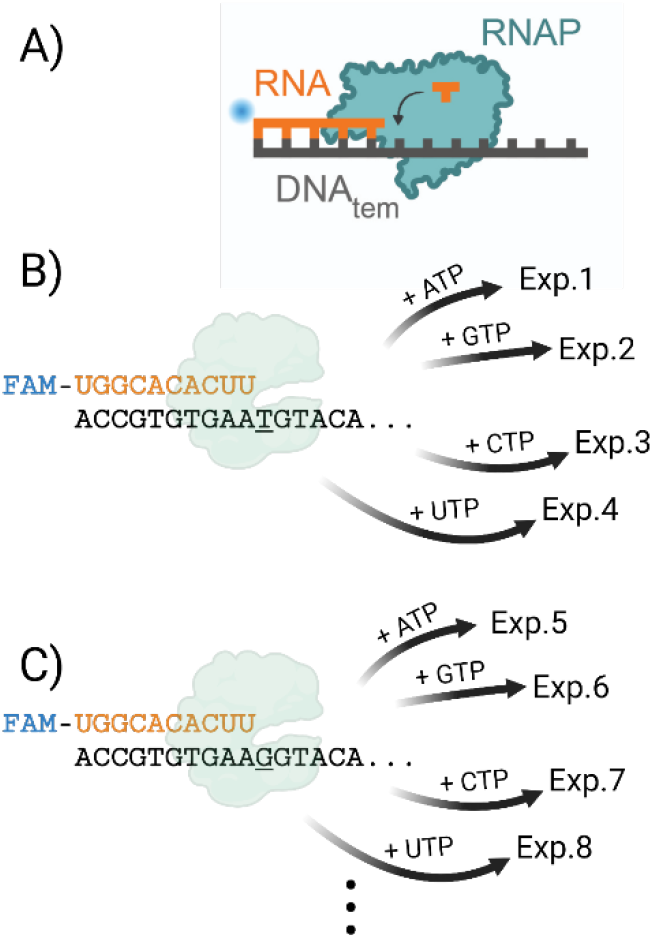
Experimental design graphic. A) Scaffold design contains a 5’ FAM labeled RNA annealed to a template DNA. RNAP is added and nucleotide addition kinetics is monitored. B) The sequence of the RNA primer is shown and a portion of the template DNA sequence is shown. The next templating nucleotide is a ‘T’. Simultaneous separate experiments are carried out where individual nucleotides are added. The first experiment (Exp.1) contains only ATP, the second experiment (Exp.2) contains only GTP, etc. C) The next group of experiments use a new template DNA which contains a templating ‘G’. Separate nucleotides are added and misincorporation kinetics is monitored. Image was generated using BioRender.

To form the elongation complex (EC), the RNA was first annealed to the single-stranded DNA template to form a DNA:RNA hybrid. Excess template DNA was used to ensure that all RNA primer was annealed. T7 RNAP was subsequently added in excess over template DNA and RNA which should result in almost all labeled RNA being incorporated into an EC (i.e. [RNAP]>[DNAtem]>[RNA]). Previous literature has shown that T7 proceeds through a slow series of conformational changes converting from an initiation complex to an elongation complex (EC) [33, 62-68]. In this platform, a 30 min pre-incubation step at 4 °C was performed to allow this slow transition to take place. The EC was then raised to the desired reaction temperature and the nucleotide addition experiment was carried out.

HiKER allows eight reactions to be carried out simultaneously and multiple sets of eight experiments to be carried out sequentially. In this experimental setup, time courses for all nucleotide combinations were monitored (*i*.*e*. rA:dG, rA:dA, rA:dC, etc.) in two sets of eight resulting in 16 unique nucleotide pairings. An illustration of the first four reactions is shown in **Fig. 3B**. The first DNA:RNA hybrid substrate has a thymine (T) templating the next RNA incorporation. The first experiment (Exp. 1), ATP is the only nucleotide in solution. Thus, the corresponding time course monitored the formation of a dT:rA. The second experiment (Exp. 2 in **Fig. 3B**) has exclusively GTP in solution. Thus, the corresponding time course monitored the formation of a dT:rG mismatch. All four ribonucleotide additions were examined by monitoring separate simultaneous experiments. The next group of four experiments were carried out where the templating “T” was exchanged for a “G” and only one ribonucleotide was in solution (**Fig.3C**). Four replicates were collected for each nucleotide pair adding redundancy to the experimental data set.

The single-subunit phage RNAPs from T3 and Sp6 were then exchanged for T7 RNAP in the above experimental setup. Time courses for each nucleotide pair were collected with four experimental replicates. In total 2304 time points (4 ribonucleotides X 4 deoxynucleotides X 3 RNA polymerases X 4 replicates X 12 time points per time course) corresponding to 192 time courses were collected using HiKER over the course of a few days.

### Overview of HiKER Platform

The HiKER (<u>Hi</u>gh-throughput <u>K</u>inetics using Capillary <u>E</u>lectrophoresis and <u>R</u>obotics) platform was developed to address the need for a high throughput kinetic system to monitor *in vitro* nucleotide addition. Currently this platform can collect ∼1500 time points a day resulting in a dramatic increase in throughput.

The first step in carrying out the misincorporation assay using HiKER was the assembly of the reaction plate (**Fig. 4A**). Each well of columns 1-3 contained individual ribonucleotides (ATP, GTP, CTP, and UTP) and columns 10-12 contained elongation complexes. A description of the EC assembly is described above and in the Materials and methods. This plate was then placed into the temperature module located within an OT-2 (Opentrons, Queens, NY) liquid handling robotics system (**Fig. 4B, Supplemental Fig.1**). Tips and time point collection plates with EDTA were also loaded into the OT-2 system to prepare for an experiment (**Supplemental Fig. 1**).

**Fig. 4.**
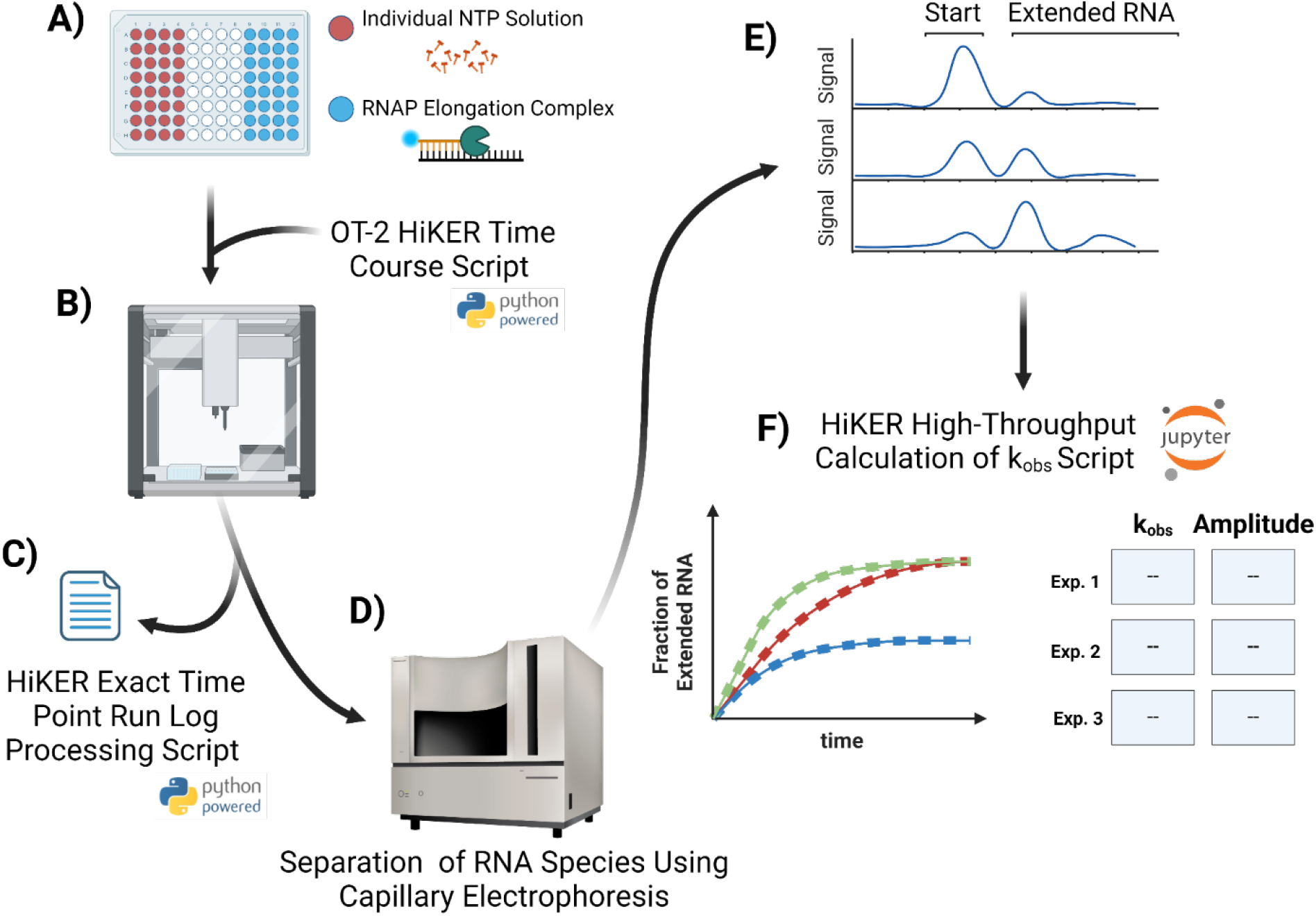
Overview of HiKER Platform. A) The reaction plate is portioned into two different sides that are mixed one to one. The left-hand side contains individual NTPs in each well (ATP, GTP, CTP, UTP). The right hand side contains elongation complexes (EC). B) The reaction plate is placed into the OT-2 system and OT-2 HiKER Time Course Script is loaded into the system. C) The HiKER Exact Time Point Run Log Processing Script is used to calculate the exact time points using the run log. D) The time point samples are loaded into a capillary electrophoresis (CE) instrument for analysis. E) The CE traces are then processed to calculate the amount of extended RNA for each collected time point. F) The HiKER High-throughput Calculation of k_obs_ Script can then ube used to visualize the time courses as well as calculate and tabulate the k_obs_ and amplitude values. This figure was generated using BioRender.

The OT-2 robotic system was chosen for the development of this platform as it is affordable and utilizes open-source scripts written in Python. We constructed a Python script which allows the OT-2 to collect time-courses and termed this script “OT-2 HiKER Time Course Script” (**Fig. 4B**). The next step in this experiment was loading this Python script into the instrument.

A caveat that should be noted is that the OT-2 (Opentrons, Queens, NY) system does not have some of the functionalities of more expensive systems; namely, it does not have a scheduling system. The scheduling feature allows a robotics system to perform movements at exact times. An example is to deliver a solution at exactly 2:32 pm. As one might expect, a kinetics experiment is extremely dependent on time. To address this challenge, a series of delay steps between time point collections was implemented. The result is a series of collected time points with defined pauses between each sample.

Upon completion of the OT-2 HiKER Time Course Script, one of the resultant outputs is a run log. The log file contains each individual step that was carried out during the execution of the script. Importantly, the delays discussed above do not account for the short amounts of time needed for the instrument to mechanically move. Consequently, the time points are not exact reaction times but instead approximate values. A separate Python script was developed to process the output log file and ascertain the exact time points adding additional rigor to this experimental approach (**Fig. 4C**).

The second output from the reaction is multiple 96 well plates containing the experimental time point samples in quenching EDTA. The samples were subsequently loaded into a capillary electrophoresis (CE) system for separation and quantification of substrates and products (**Fig. 4D**). CE is a technique that is similar to PAGE where mixtures of different components are separated based on size and electrostatic properties using a polymer matrix [69]. CE is an optics-based system where a dye must be present on the RNA/DNA for detection.

This methodology has shown potential in a variety of applications [61, 70-79]. The 3730 DNA Analyzer from ThermoFisher Scientific (Waltham, MA) was used in this study and processes 96 samples simultaneously. The system is extremely efficient using 1 µL of sample and requiring approximately one hour of run time. The 3730 DNA Analyzer is the largest capital expense of the systems but more affordable options are also available. The output from this system are individual CE traces for each sample (**Fig.4E)**. The fraction of extended RNA was calculated for each time point and time courses were constructed (**Fig. 4F**). A final Python script was developed to aid in processing the large volume of time courses. The time courses are fit to a single exponential (Eq. 1) and the resultant k_obs_ and amplitude parameters are tabulated.

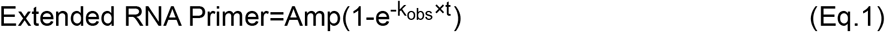

### T7 RNAP Misincorporation Kinetics

Using HiKER, we first probed T7 RNAP misincorporation kinetic profiles and compared the observations with previously published findings. **Fig. 5A** contains time courses for each of the four ribonucleotides with a deoxyadenosine (A) on the template DNA strand. The reaction was carried out in single-turnover conditions ([E]>>[S]) and the nucleotide concentration was held in excess. As expected, addition of correctly paired dA:rU was fastest with the reaction reaching completion in under 15 s. The reaction did not go to 100%, but rather plateaued at ∼75%. The incomplete reaction has been observed in previous publications and is suggested to be a consequence of arrested ECs [32, 33]. Three or more experimental replicates were collected and the average time courses were fit using Eq. 1 and the data analysis tool KaleidaGraph (Synergy Software, Reading, PA). Two optimized parameters were determined for each fit, an amplitude (Amp) and an observed rate-constant (k_obs_). The resultant fit parameters were tabulated and are shown in **Supplemental Table 1**.

**Table 1.**
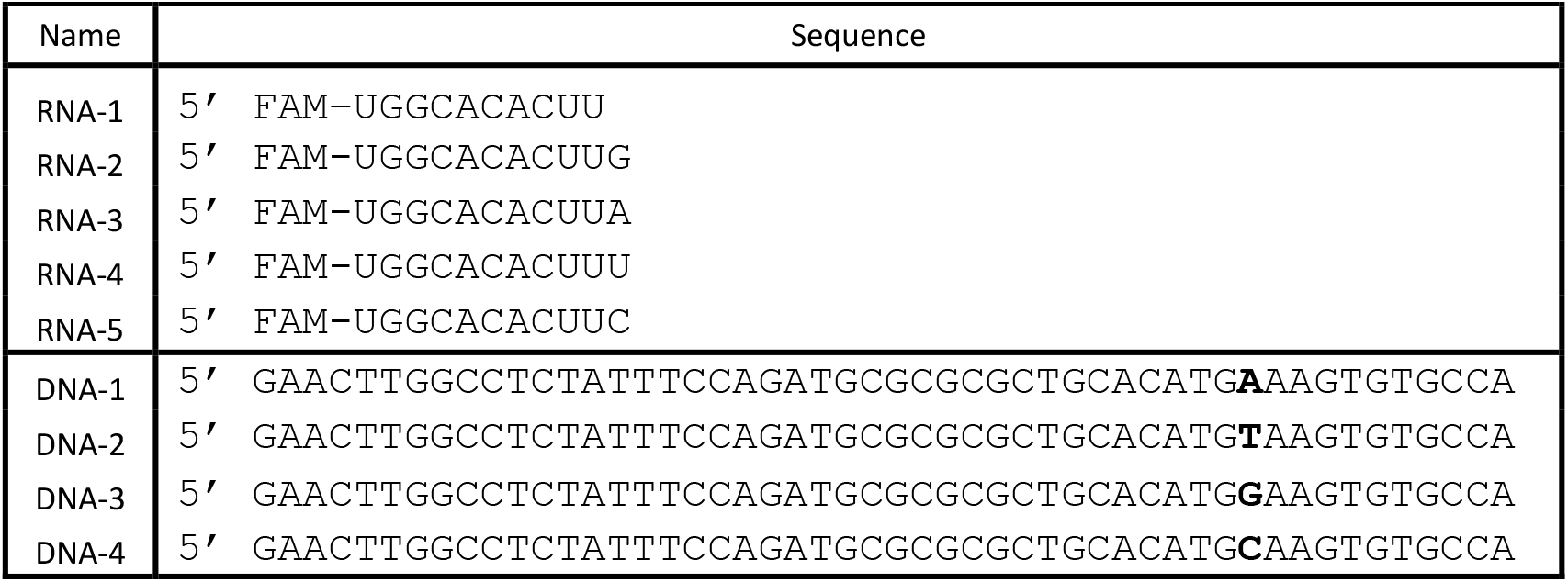
DNA and RNA Sequences.

**Fig. 5.**
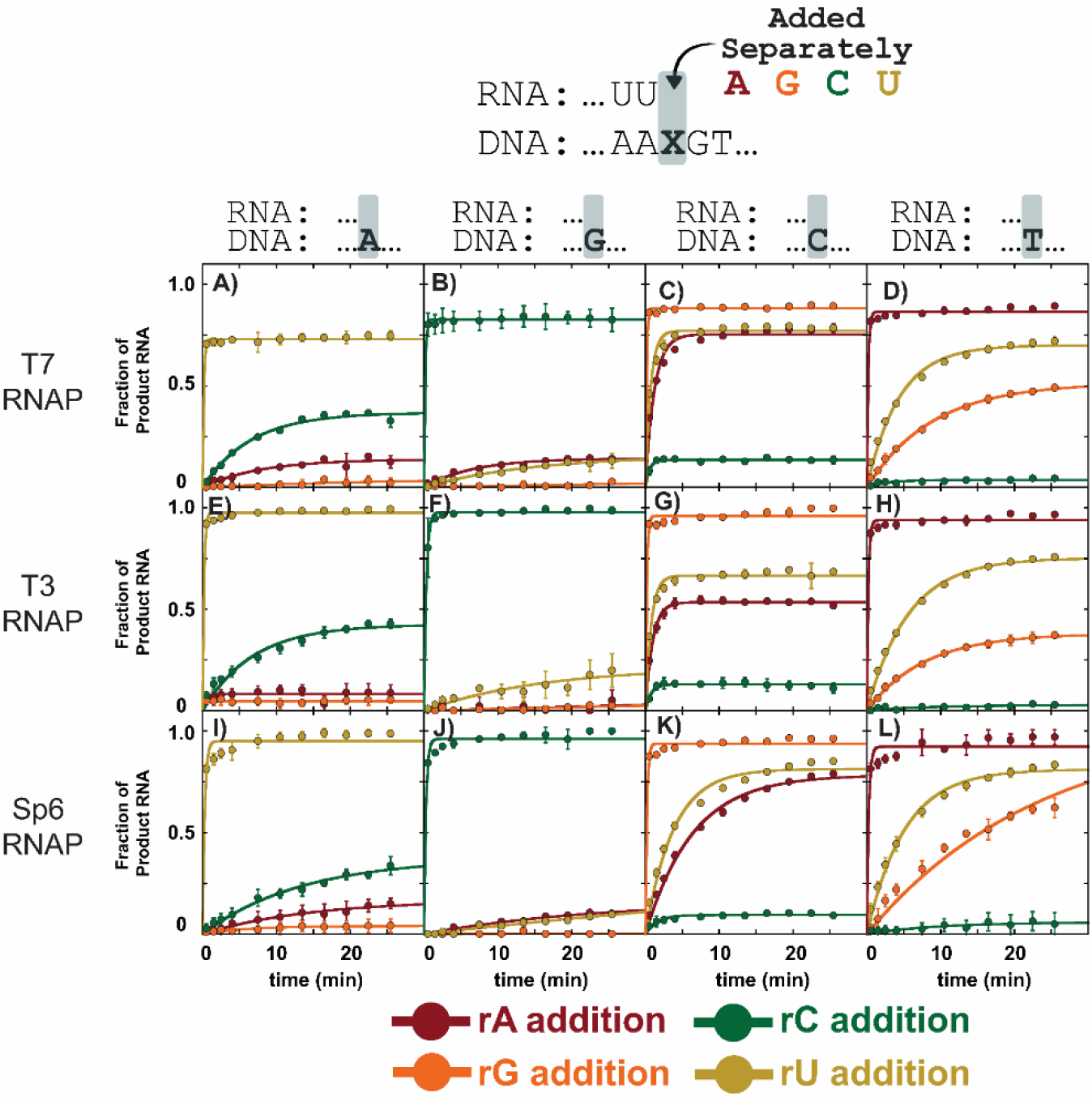
Misincorporation kinetics for WT RNAP enzymes T7, T3, and Sp6 RNAP in a variety of sequence contexts using HiKER. Each row is a different RNA polymerase, each column is a different nucleotide on the template strand, and each color time course corresponds to a different ribonucleotide incorporation. Experiments were carried out in single-turnover conditions with 1 mM of the NTP in solution. Experiments were carried out with at least three experimental replicates. The data points are the average, and the error bars are the standard deviations. Additional experimental conditions can be found in the Materials and Methods.

Upon inspection of the time courses in **Fig. 5A**, an immediate observation is the differences in amplitude. rC addition had the largest amplitude of the misincorporated nucleotides followed by rA and rG. Both rU and rC are pyrimidines. This observation indicates T7 RNAP more readily forms a purine-pyrimidine mismatch compared to a purine-purine mismatch. This selectivity is consistent with previous misincorporation studies on T7 RNAP [7, 35] as well as *E. coli* RNAP [47]. Interestingly, the k_obs_ for dA:rC is only marginally faster when compared to the k_obs_ for dA:rA. These experiments were carried out simultaneously and under identical experimental conditions suggesting clear mechanistic differences between the two mismatches.

Time-courses for the remaining 12 correct and incorrect nucleotide pairings were carried out (**Fig. 2 B-D**). As expected, the correct ribonucleotide base for each template deoxynucleotide was incorporated fastest and the reaction was complete by 15 s. The dG template exhibited the greatest specificity with minimal misincorporation (**Fig. 2B**). In contrast, the dC template had both the fastest observed rate-constants (∼1.8 min^-1^) for misincorporation as well as the highest amplitudes (**Fig. 2C)**. The tabulated observed rate-constants are shown in **Supplemental Table1**. Interestingly, rG misincorporation by T7 RNAP appeared to be an unfavorable reaction with no significant amount of misincorporation at templating dA or dG positions (**Fig. 5A, 5B**). Misincorporation of rG only appeared at templating dT position (**Fig. 5D**).

### Comparing Misincorporation Kinetics of T7, T3, and Sp6 RNAP

The three bacteriophages T7, T3, and Sp6 each contain a ∼99 kDa single subunit RNAP. The structure function relationship suggests that T7, T3, and Sp6 would have similar mechanistic properties [80, 81]. However, it should be noted that opposing phenotypes have been observed in structurally similar enzymes [82-84]. Thus, structural similarities between enzymes ‘suggests’ similar chemical properties, but in solution experimental evidence is needed to test this hypothesis. With this high-throughput platform in hand, we sought to directly compare the misincorporation profiles of T7, T3, and Sp6 RNAP under identical experimental conditions.

Using HiKER, 16 different time courses encompassing the four ribonucleotide possibilities on the RNA strand and the four deoxynucleotides on the template DNA strand were collected for T7, T3, and Sp6 RNAP (**Fig.5**). Each time course was fit using Eq.1 and the resultant fit parameters are tabulated in **Supplementary Table 1**. Generally, the three enzymes exhibited similar misincorporation properties. Strikingly, misincorporation appears less favorable at template purines positions (first two columns of **Fig. 5**) compared to template pyrimidine positions (last two columns of **Fig. 5**). The changes in both k_obs_ and amplitude throughout suggest the selection of the correct nucleotide is not simply a consequence of being kinetically favorable.

It should be highlighted that studies using sequencing gels are often forced to focus on a relatively small subset of mismatch formations and then extrapolate the findings to make broad conclusions. With HiKER all misincorporations were monitored and the unique features for individual mismatches were observed.

### T7, T3, and Sp6 RNAP extension from a mismatch

Two key steps must occur for the existence of an error in a synthesized RNA transcript. First, the incorrect nucleotide must be incorporated into the nascent RNA strand, and then the RNAP must extend from a base pair mismatch. Here we utilized HiKER to investigate the reaction kinetics for T7, T3, and Sp6 RNAP extension from all base combinations. RNA primers with different 3’ terminal nucleotides were annealed to different template DNAs to produce all 16 possible combinations of pairings between DNA and RNA. All four nucleotides were added and the amount of extension was monitored (**Fig. 6**).

**Fig. 6.**
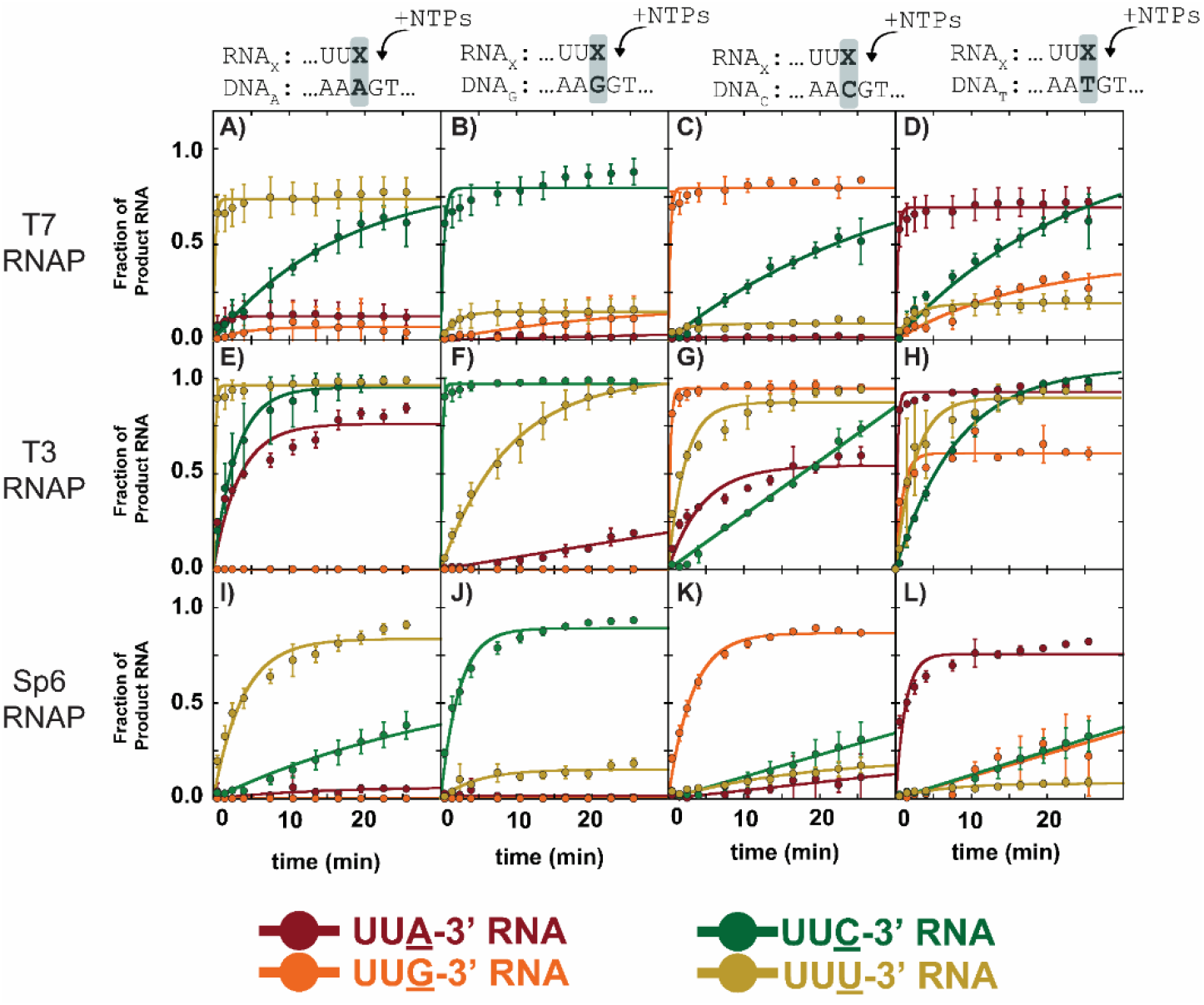
T7 RNAP extension from mismatch kinetics using HiKER platform. A-L) Each panel corresponds to one of the four deoxynucleotides on the template DNA. Within each panel there are four time courses corresponding to four different RNA primers with different 3’ ends. The different combinations result in different mismatches. 1 mM of each NTP was added to each elongation complex to measure extension from a different DNA:RNA pair. Experiments were carried out under single-turnover conditions. Experiments were carried out with at least three experimental replicates. The data points are the average, and the error bars are the standard deviations. Best fit lines correspond to fits using a single exponential. Additional experimental details can be found in the materials and methods.

The first set of experiments monitored extensions from the mismatches dA:rA, dA:rG, and dA:rC (**Fig. 6A**). Additionally, extension from correctly paired dA:rU was observed. As expected, the correctly matched RNA primer (…UUU-3’) extended the fastest with the reaction reaching completion prior to the first reaction time point at 15 s. ∼75% of the RNA primer was extended, consistent with observations observed in **Fig. 5A**.

Much like in the previous set of experiments, T7 RNAP extension from 16 different mismatch combinations was explored (**Fig. 5 A-D**). The time courses were fit using a single exponential (Eq.1) and the resultant fit parameters are tabulated in Supplemental Table 2. The low amplitude and slow reaction kinetics indicate T7 RNAP does not extend well from a mismatch and is consistent with previous studies [7, 35]. Interestingly, extensions from purine:purine mismatches resulted in time courses with very low amplitudes (**Fig. 6**) suggesting purine-purine mismatches are extremely unfavorable. Mismatches at a templating G (**Fig. 6B**) showed the least amount of extension from a mismatch.

Time courses monitoring the extension from different mismatches by T3 and Sp6 RNAP were also collected (**Fig. 6 E-L**). In contrast to the misincorporation formation kinetics shown in **Fig. 5**, extension from a mismatch kinetics were dramatically different between the RNA polymerases. Consider the first column of **Fig. 6** which corresponds to mismatches including a dA on the template DNA. The extension from correct dA:rU is fast and with a high amplitude for both T7 and T3 RNAP (**Fig. 6A** and **Fig. 6I**). Strikingly, for Sp6 RNAP the dA:rU extension was dramatically slower. Two potential explanations for this observation are that Sp6 is bound but not able to rapidly extend from this DNA:RNA construct or Sp6 is dissociating from this construct and the observed signal is a combination of rebinding steps followed by nucleotide addition. Differences in extension from a mismatch at templating A were also observed between T7, T3, and Sp6 RNAP (**Fig. 6A, 6E, and 6I**). The trend was consistent with dA:rC being the fastest and most abundant followed by dA:rA and no extension at dA:rG.

Using the HiKER platform, extension from other base combinations was explored across T7, T3, and Sp6 RNAP (**Fig. 6**). Strikingly, differences were observed between the polymerases. T3 RNAP more readily extended from a mismatch compared to T7 and Sp6 RNAP. Extension from mismatches with a dG was minimal, with the exception of dG:rU by T3 RNAP (**Fig. 6F**). Extension from the correctly matched base pairs was slow in all cases for Sp6 RNAP, but fast for T7 and T3 RNAP. Extension of mismatches appears slower for Sp6 compared to T7 and T3 RNAP.

## Discussion

### Changes in Amplitude for Different Mismatches

The changes in amplitude for different misincorporations is an intriguing observation (**Fig. 5**). While T7 RNAP has been well studied, little mechanistic information is known about T3 and Sp6 RNAP. The similarities between RNAPs observed in **Fig. 5** suggests a conserved kinetic mechanism for misincorporation between the polymerases but additional future experiments are needed to explore this further.

It has previously been reported that the T7 RNAP Kd for pyrophosphate is ∼1.2 mM and pyrophosphate release is fast relative to other kinetic steps in correct T7 RNAP nucleotide addition [33]. The free pyrophosphate in solution for the misincorporation experiments would be extremely low, thus binding of free pyrophosphate is unlikely. Thus, for the pyrophosphorolysis step (reverse of polymerization step) to be present, the pyrophosphate from the incoming NTP could not be released. One hypothesis is that pyrophosphate release is slow for misincorporation, but an alternative hypothesis is that bond formation is slow for misincorporated nucleotides and of similar magnitude to pyrophosphorolysis at a mismatch. Pyrophosphorolysis by *E. coli* RNAP has been reported to occur on the time scale of minutes[85]. The observed changes in amplitude would occur if pyrophosphorolysis by T7 RNAP is occurring on a similar time scale and misincorporation.

We used simulations to take first steps in identifying a mechanism that would give rise to changes in the amplitude shown in **Fig. 5**. Four different reaction schemes are shown in **Fig. 7** with corresponding simulated time courses. One kinetic step was varied in each case with the goal to reproduce the observed differences in amplitude. The rationale for each mechanism shown in **Fig. 7** is as follows: Scheme 1 evaluates the impact of a reversible bond formation step, Scheme 2 evaluates including an open to closed conformation change prior to the bond formation step, Scheme 3 tests the impact of the elongation complex existing in a pre-equilibrium prior to nucleotide binding, and lastly Scheme 4 shows the impact an off pathway conformational change after nucleotide binding. Fits using a single exponential (**Eq.1**) were performed to each simulated time course and the resultant fit amplitude are shown in **Fig. 7**. From inspection of the simulated time courses and the changes in the fit amplitude Scheme 1 and Scheme 4 appear to best describe the experimental data shown in **Fig. 5**. However, additional mechanistic studies are needed to elucidate the mechanism of nucleotide misincorporation.

**Fig. 7.**
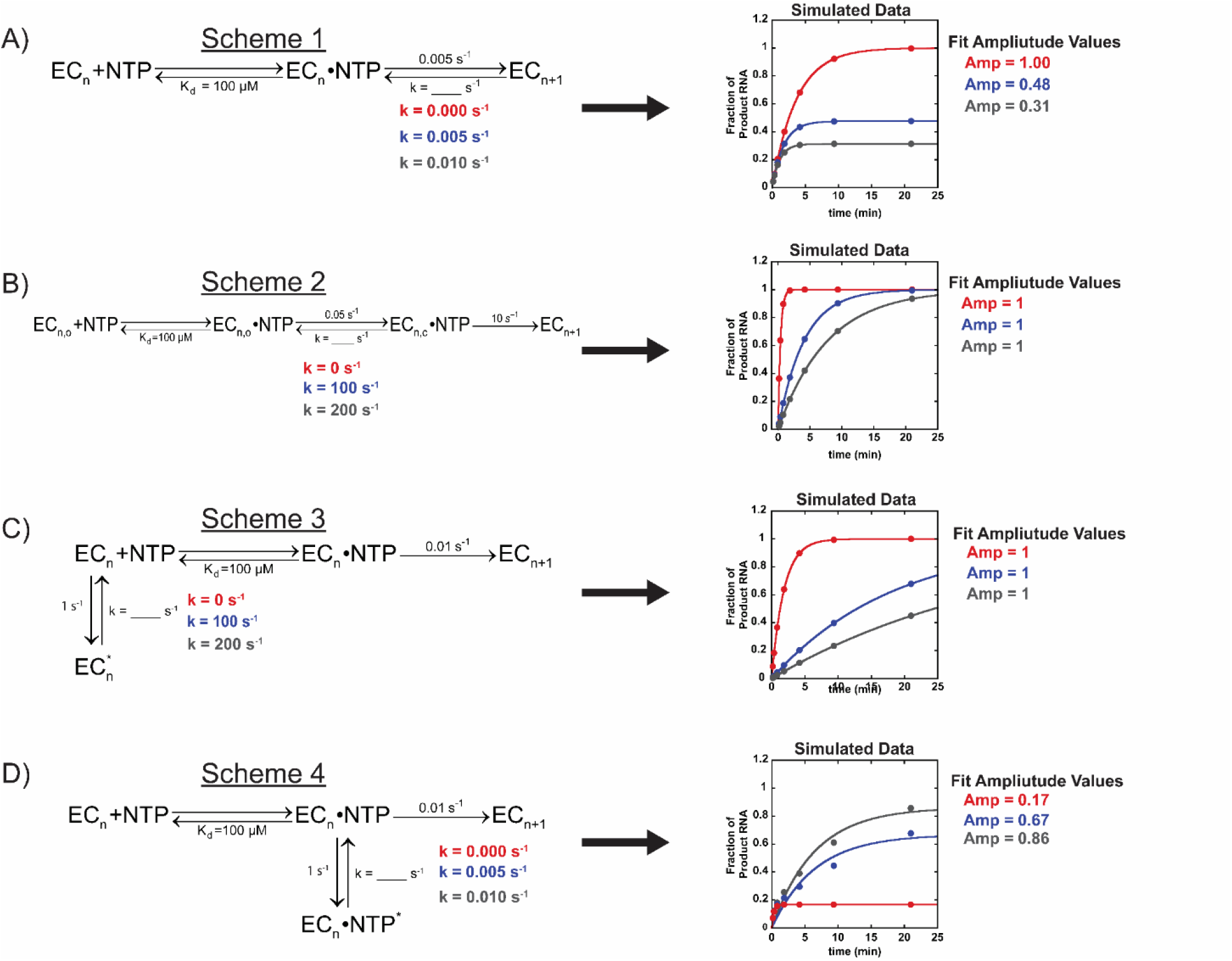
Simulations probing different kinetic mechanisms as well as investigating the impact of individual rate constant values on final amplitude. A) Simulation where bond formation step is reversible. B) Simulation including an open and closed conformational change prior to an irreversible bond formation step. C) Simulation constructed where elongation complex exists in equilibrium between active and inactive state prior to nucleotide binding. D) Simulation where a portion of the elongation complexes go through a conformational change prior to bond formation. Simulations were made using MENOTR and fit to Eq.1 using KaleidaGraph. Additional details can be found in the materials and methods.

In previous literature studies, amplitude has often been attributed to a measure of active elongation complexes and has been largely ignored. However, in this high-throughput study, identical elongation complexes were assembled in the same tube with minimal differences between experiments. This is evident by the small error bars shown in **Fig. 5**. The changes in amplitude are thus a reflection of mechanistic differences in nucleotide addition rather than differences in sample preparation.

Molecular dynamics simulations of T7 nucleotide addition indicate that the correct nucleotide forms preferable contacts with key residues and can be well-stabilized [86, 87]. In contrast, incorrect substrates exhibit greater flexibility in the insertion site. The changes in amplitude could reflect differences in mismatch stability. Overall, mismatches across a dG exhibited lower amplitudes (∼< 0.1) while mismatches across from dC exhibited some of the largest (∼0.6). This suggests that mismatches at dG are the least stable and mismatches at dC are the most stable.

### Misincorporation frequency predictions from branching ratio

Previous sequencing results indicate T7 misincorporates ∼46 misincorporations per million bases [5, 6]. How do the T7 RNAP misincorporation results presented here using the high-throughput kinetics platform compare? The ratio of the observed rate constant for misincorporation and the observed rate constant for correct nucleotide addition provides an estimate of the likelihood for an individual misincorporation. The correct nucleotide addition was too rapid to capture using this assay, but previous studies report a k_obs_ value of ∼200 s^-1^ under saturating NTP conditions [34, 88-90]. Therefore, the mean of the ratio of incorrect versus correct nucleotide addition across multiple sequence contexts is ∼45 misincorporations per million bases.

The consistency between calculated misincorporation frequencies using either next generation sequencing or HiKER is striking as the experimental strategies are vastly different. Misincorporation from sequencing studies reflects many different sequence contexts while the HiKER platform measured the k_obs_ for nine different mismatch formations with consistent upstream and downstream template sequences. Previous studies have shown differences in misincorporation kinetics at different templating positions suggesting sequence effects [7]. While this question was outside the scope of this study, HiKER allows for the rapid investigation of many different sequence contexts thus serving as a powerful tool.

A similar prediction of the misincorporation frequency can be calculated for T3 RNAP. The average misincorporation k_obs_ observed here is ∼0.78 min^-1^ and the correct nucleotide addition k_obs_ has been estimated to be ∼170 s^-1^ [60]. This ratio suggests a misincorporation frequency of ∼76 misincorporations per million bases. This value is similar to the previously value of ∼48 misincorporations per million bases from NGS sequencing experiments [5].

Sp6 RNAP is the least characterized of the RNAPs investigated here. To our knowledge the k_obs_ for correct nucleotide addition by Sp6 RNAP has not been reported. If we assume the value is comparable to other phage RNAPs the value should be ∼185 s^-1^ (average between T7 and T3 values). Using this value the estimated error frequency is ∼21 misincorporations per million bases. Interestingly, the published NGS sequencing results report a value of ∼118 misincorporations per million bases. The disparity between these two values is potentially the result of an incorrect k_obs_ value for correct nucleotide addition. A recalculation of the k_obs_ for correct nucleotide addition using the sequencing misincorporation frequency value of ∼110 misincorporations per million bases and the average misincorporation k_obs_ value of ∼0.23 min^-1^ reported here gives a predicted value of ∼33 s^-1^. This suggests that correct nucleotide addition by Sp6 RNAP is much slower compared to T7 and T3 RNAP, but more comparable to multi-subunit RNAP’s found in *Escherichia coli* and eukaryotes [38, 40, 41, 48, 50, 54].

### HiKER Platform empowers high-throughput enzymology

Despite many technological advances, PAGE gels have remained the gold standard for many nucleotide addition studies. Advances towards high-throughput approaches in binding affinity measurements using surfaced plasmon resonance (SPR) and biosensor technologies have been developed [91, 92]. Custom devices have been developed using microfluidics and allow hundreds to thousands of reactions to be investigated in short time spans [93]. However, all these technologies require substantial capital investment by researchers. Recent publications have demonstrated that affordable liquid handling robotic systems can be used to carry out kinetic experiments monitoring organic and inorganic chemical reactions [94, 95]. HiKER is a step towards empowering enzymologists and biophysicists to carry out high-throughput biochemical reactions. While HiKER was setup for CE, the platform is easily adapted to use other analytical methods such as mass-spectrometry or HPCL.

All experimental studies have limitations and drawbacks. A current drawback for this system is that it is unclear if the observations reflect actively transcribing ECs. Preliminary data indicate HiKER is consistent with previous literature, but additional controls and modifications must inevitably be made to better correlate with transcribing ECs. One key benefit of using a high-throughput system is the ability to directly compare performance in a wide variety of different experimental conditions. The *in vitro* experimental setup may not directly mimic cellular conditions, but the trends in the enzymatic properties are likely consistent. The observations presented in this study demonstrate the capabilities of HiKER. This platform readily enables the optimization of incorporation of modified nucleotides, important in RNA therapeutics and vaccines, as well as the optimization of reaction conditions and high throughput testing of new RNAP mutants and homologs.

### Conclusion

Here we have developed a high-throughput kinetics platform that allows many enzymatic reactions to be monitored simultaneously. We leveraged this technology to explore the misincorporation kinetics of T7, T3 and Sp6 RNAP and demonstrated similar misincorporation properties. We then explored how RNAPs extend from a mismatch and observed differences between the RNAPs with T3 RNAP most readily extending from a mismatch and Sp6 RNAP extending the least. The experiments comparing different RNAPs presented here were carried out with identical experimental conditions allowing direct comparisons without the potential complications of comparing different studies and experimental designs. Predicted misincorporation frequencies in a full-length RNA transcript were comparable to reported values using HiKER. The results presented here demonstrate the capabilities of HiKER and serve as a critical tool for future studies by providing a framework for future studies to enable high-throughput characterization of many different RNA/DNA polymerases as well as being adapted for other enzyme systems.

## Materials and methods

### Reagents

All enzymes and reagents used in this study were from New England Biolabs (Ipswich, MA, USA) unless otherwise stated. T7 RNAP, T3 RNAP, and Sp6 stocks were prepared, and the concentrations were determined to be 176 μM, 48 μM, and 102 μM respectively. The reactions shown in **Fig. 5** and **Fig. 6** were carried out in RNAP Buffer. The composition of this buffer is 40 mM Tris-HCl (pH 7.9 @ 25°C), 6 mM MgCl2, 1 mM DTT, 2 mM spermidine.

The synthetic oligonucleotides used in this study were purchased from Integrated DNA Technologies (Coralville, IA). RNase free HPLC purification was performed for each RNA and DNA oligo. **Table 1**. contains the oligo sequences.

### Misincorporation Assay and Extension from Mismatch Assay Using HiKER

This kinetic assay was initiated by mixing two 50 µL mixtures 1:1. The two mixtures were the EC mixture and the NTP mixture. The EC mixture was composed of 1X RNAP Buffer, 0.3 µM RNA primer, 1 µM template DNA. This was distributed in 50 uL aliquots to the right hand columns of a 96 well plate (**Supplemental Fig. 1**). The DNA:RNA hybrid was formed with the use of a thermocycler. The annealing protocol is to heat to 95 °C and then drop 0.1 °C per second until 4 °C is reached. Titrations of increasing [RNA Polymerase] were performed until no change in reaction kinetics was observed suggesting single-turnover conditions being achieved. The volume of the RNAP that was added for three polymerases is as follows: 1.5 µL of 176 µM T7 RNAP, 3 µL of 48 µM T3 RNAP, 1.5 µL of 102 µM Sp6 RNAP.

The EC mixture is then incubated for 30 min at 4 °C and subsequently raised to the desired reaction temperature for 5 min. During this incubation time 100 uL of individual NTP’s were loaded into the right hand columns of the reaction plate (**Supplemental Fig. 1**). The reagent plate was then loaded into the Opentrons (Queens, NY) temperature module (Gen2) located within the OT-2 liquid handling robot.

Sample collection plates were pre-filled with 10 uL of 50 mM EDTA to quench the reaction and placed with tips into the OT-2 (**Supplemental Fig. 1**).

### Time-course collections using OT-2

The OT-2 HiKER Time Course Script and all HiKER scripts are located in the **Supplementary data**. The script is loaded into the instrument and a series of steps take place. Step 1: 2 μL of the first column of the EC samples is transferred to the first column of the sample collection plate and corresponds to zero time point. Step 2: 50 uL of the first column of the NTP samples is transferred to the first column of the EC samples and the solution is mixed. Step 3: 10 uL of the mixed solution is transferred to the sample collection plate and is the second time point (zero being the first). This process is repeated until all 12 time points are collected filling a 96 well plate. Step 4: This process is repeated until all reactions are completed (4 times max – See Supplemental Fig.1). Once complete. The samples are diluted with 100 uL of H20.

### Capillary Electrophoresis Run Details

An 3730xl DNA Analyzer from Applied Biosystems (Waltham, MA) was used to perform the capillary electrophoresis experiments shown in this manuscript. The polymer POP-6 for 3730/3730xl DNA Analyzer from Applied Biosystems (Waltham, MA) was used in the instrument. Running of the samples began with 15 μl of GeneScan 120 LIZ dye size standard being added to 1000 μl of Hi-Di Formamide. Hi-Di Formamide is used to stabilize denatured DNA samples before performing CE. 10 uL of the LIZ/Hi-Di mixture was added to each sample in the 96-well CE plate. 1 μL of each experimental sample is added to the corresponding well in the CE plate. The samples are spun down, a septa is placed on top, and then loaded into a cartridge for running.

### Data Analysis and Simulations

A Python script was developed to extract out the exact time point using the run log. The name of the python script is “HiKER Exact Time Point Run Log Processing Script”. A copy of the python script is located in the **Supplementary Data**. Area under peaks was quantified using PeakScanner (ThermoFisher Scientific, Waltham, MA). Data analysis was performed using KaleidaGraph (Synergy Software, Reading, PA). A Jupyter notebook was developed to aid with analyzing the data. A copy of the script is in the Supplementary Data. The script is designed to be used with a single 96 well plate where 8 time courses are collected simultaneously. The script fits the data using Eq. 1 and tabulates the best fit parameters in a separate excel document. Simulated time courses were generated using MENOTR [96] and MATLAB (MathWorks, Natick, MA). The set of ordinary differential equations corresponding to each scheme are as follows.

#### Scheme 1 Differential Equations

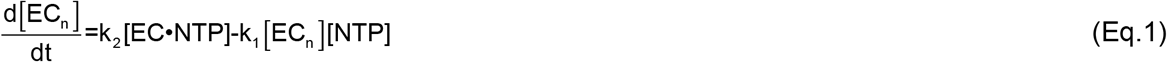

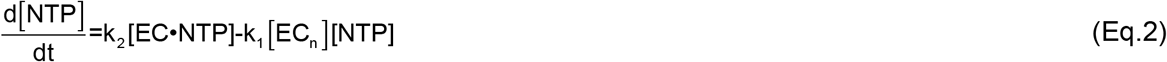

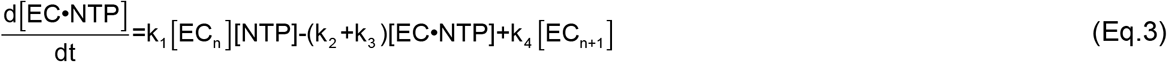

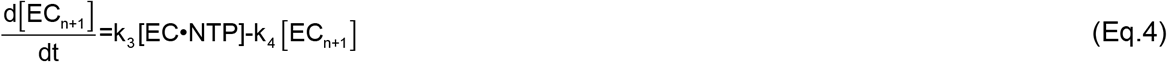

#### Scheme 2 Differential Equaitons

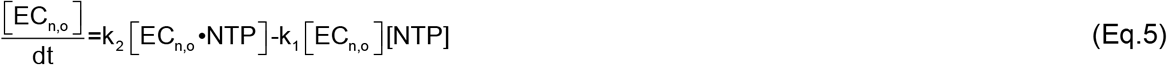

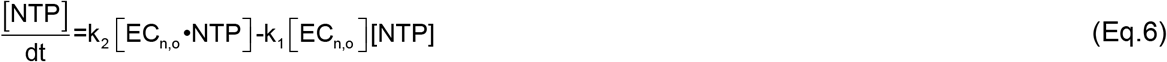

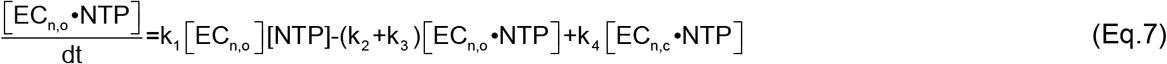

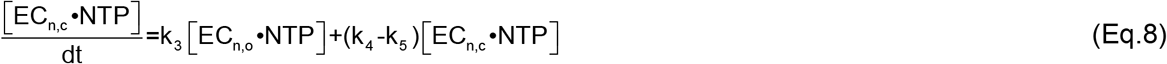

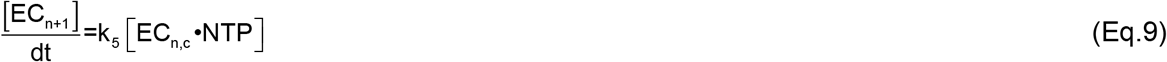

#### Scheme 3 Differential Equations

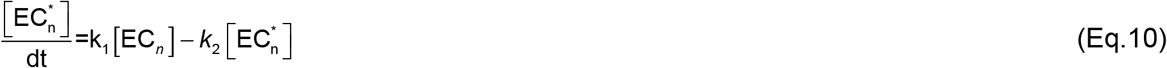

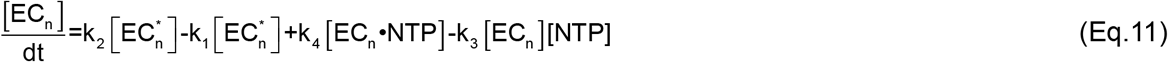

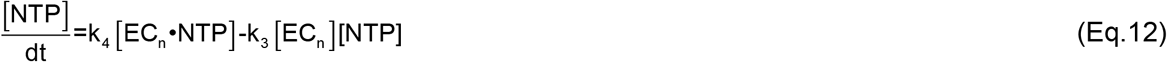

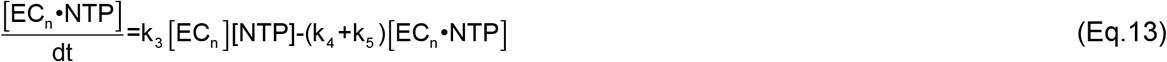

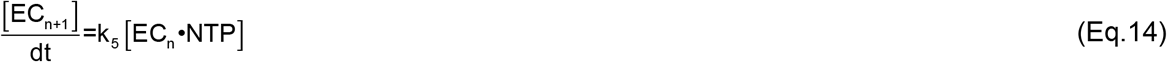

#### Scheme 4 Differential Equations

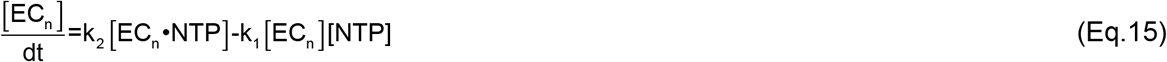

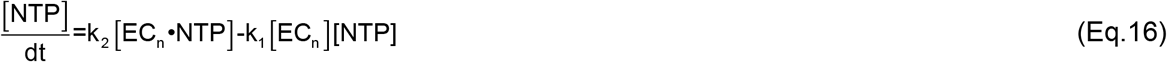

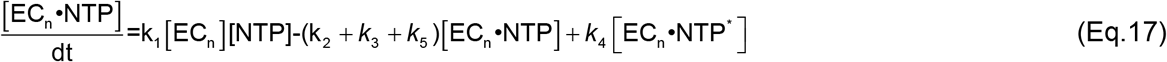

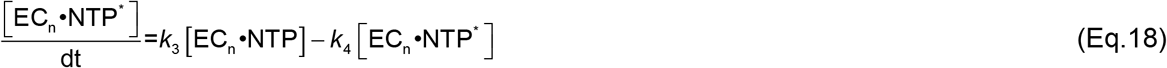

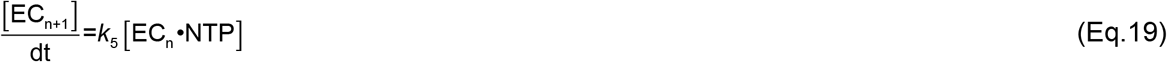

## Acknowledgements

The authors would like to thank Kelly Zatopek, Bill Jack, Christa Molé, Tien-Hao Chen, Juan Pan, Lana Saleh, members of the and the NEB Research Department for providing feedback throughout this project. The authors would like to thank Tasha José for her work on the HiKER logo design and the NEB Sequencing Core for running the numerous capillary electrophoresis samples.

