## Supplemental Document for "High-throughput Kinetics using Capillary Electrophoresis and Robotics (HiKER) platform used to Study T7, T3, and Sp6 RNA Polymerase Misincorporation"

#### Supplemental Fig. 1

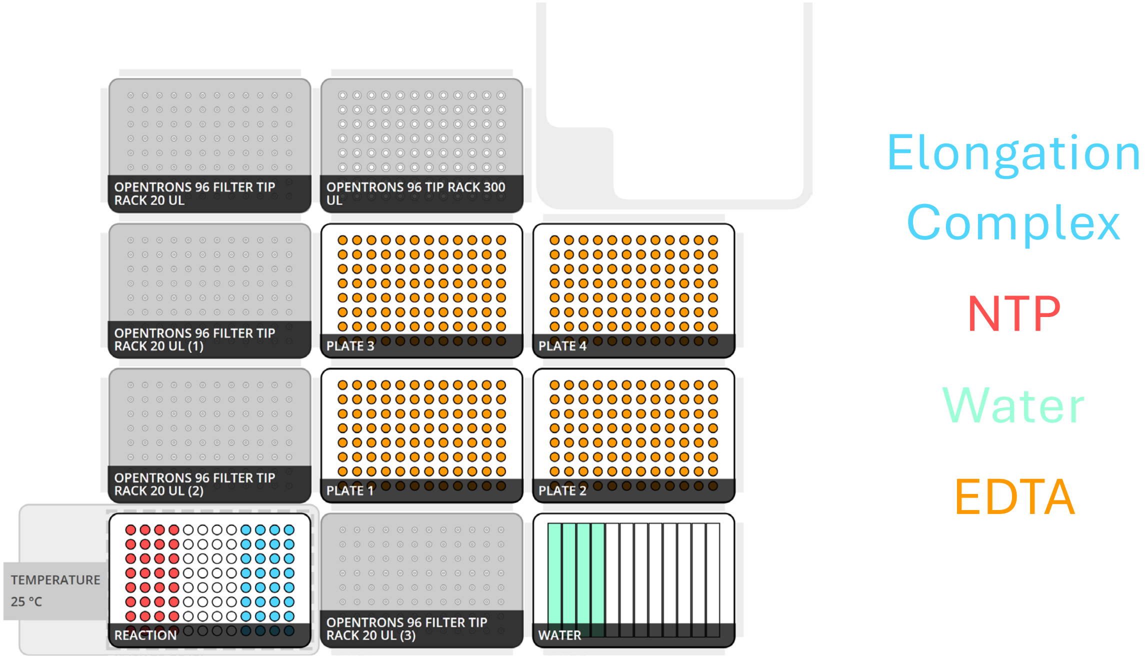


General reagent layout for OT-2 deck when using HiKER to perform nucleotide addition assays. Plate 1 – Plate 4 contains sample collection plates containing excess EDTA to quench the reaction. The reaction plate contains two different groups of reagents. The red left hand side contains the individual NTP solutions in separate wells. The blue right hand side group contains the elongation complex scaffolds.

### Supplemental Table 1

| Supplemental Table 1: Misincorporation Formation Kinetics of T7, T3, and Sp6 RNA Polymerase | | | | | | | | |
|---|---|---|---|---|---|---|---|---|
| T7 RNAP | | | T3 RNAP | | | Sp6 RNAP | | |
| | $k_{obs}$ ($min^{-1}$) | Amp | | $k_{obs}$ ($min^{-1}$) | Amp | | $k_{obs}$ ($min^{-1}$) | Amp |
| **dA:rA** | 0.13 ± 0.03 | 0.13 ± 0.01 | **dA:rA** | 2 ± 1 | 0.083 ± 0.007 | **dA:rA** | 0.07 ± 0.02 | 0.17 ± 0.02 |
| **dA:rG** | ND | ND | **dA:rG** | ND | ND | **dA:rG** | 0.17 ± 0.05 | 0.04 ± 0.01 |
| **dA:rC** | 0.15 ± 0.01 | 0.37 ± 0.01 | **dA:rC** | 0.15 ± 0.02 | 0.42± 0.02 | **dA:rC** | 0.07 ± 0.01 | 0.37 ± 0.03 |
| **dA:rU** | > 5 | 0.73 ± 0.01 | **dA:rU** | > 5 | 0.97 ± 0.01 | **dA:rU** | > 5 | 0.95 ± 0.01 |
| **dG:rA** | 0.15 ± 0.02 | 0.14 ± 0.01 | **dG:rA** | ND | ND | **dG:rA** | 0.04 ± 0.02 | 0.17 ± 0.05 |
| **dG:rG** | ND | ND | **dG:rG** | ND | ND | **dG:rG** | ND | ND |
| **dG:rC** | > 5 | 0.83 ± 0.01 | **dG:rC** | > 5 | 0.98 ± 0.01 | **dG:rC** | > 5 | 0.96 ± 0.01 |
| **dG:rU** | 0.07 ± 0.01 | 0.16 ± 0.01 | **dG:rU** | 0.06 ± 0.03 | 0.22 ± 0.06 | **dG:rU** | 0.001 ± 0.02 | 0.5 ± 0.9 |
| **dC:rA** | 0.82 ± 0.09 | 0.75 ± 0.01 | **dC:rA** | 1.03 ± 0.05 | 0.53 ± 0.01 | **dC:rA** | 0.78 ± 0.02 | 0.16 ± 0.01 |
| **dC:rG** | > 5 | 0.88 ± 0.01 | **dC:rG** | > 5 | 0.96 ± 0.01 | **dC:rG** | > 5 | 0.94 ± 0.01 |
| **dC:rC** | 1.8 ± 0.2 | 0.14 ± 0.01 | **dC:rC** | 1.4 ± 0.3 | 0.13 ± 0.01 | **dC:rC** | 0.6 ± 0.1 | 0.09 ± 0.01 |
| **dC:rU** | 1.4 ± 0.2 | 0.77 ± 0.01 | **dC:rU** | 1.3 ± 0.1 | 0.67 ± 0.01 | **dC:rU** | 0.26 ± 0.02 | 0.81 ± 0.02 |
| **dT:rA** | > 5 | 0.86 ± 0.01 | **dT:rA** | > 5 | 0.94 ± 0.01 | **dT:rA** | > 5 | 0.92 ± 0.02 |
| **dT:rG** | 0.11 ± 0.01 | 0.51 ± 0.01 | **dT:rG** | 0.13 ± 0.01 | 0.38 ± 0.01 | **dT:rG** | 0.04 ± 0.01 | 1.0 ± 0.2 |
| **dT:rC** | ND | ND | **dT:rC** | ND | ND | **dT:rC** | 0.10 ± 0.05 | 0.05 ± 0.01 |
| **dT:rU** | 0.23 ± 0.02 | 0.70 ± 0.01 | **dT:rU** | 0.18 ± 0.01 | 0.75 ± 0.01 | **dT:rU** | 0.20 ± 0.01 | 0.81 ± 0.01 |

### Supplemental Table 2

| Supplemental Table 2: T7, T3, and Sp6 RNA Polymerase Extension from a Mismatch Kinetics | | | | | | | | |
|---|---|---|---|---|---|---|---|---|
| T7 RNAP | | | T3 RNAP | | | Sp6 RNAP | | |
| | $k_{obs}$ ($min^{-1}$) | Amp | | $k_{obs}$ ($min^{-1}$) | Amp | | $k_{obs}$ ($min^{-1}$) | Amp |
| **dA:rA** | 1.3 ± 0.2 | 0.13 ± 0.01 | **dA:rA** | 0.29 ± 0.06 | 0.76 ± 0.04 | **dA:rA** | 0.07 ± 0.07 | 0.06 ± 0.03 |
| **dA:rG** | 0.2 ± 0.1 | 0.07 ± 0.01 | **dA:rG** | ND | ND | **dA:rG** | ND | ND |
| **dA:rC** | 0.06 ± 0.01 | 0.84 ± 0.08 | **dA:rC** | 0.34 ± 0.02 | 0.95 ± 0.02 | **dA:rC** | 0.03 ± 0.01 | 0.7 ± 0.3 |
| **dA:rU** | > 5 | 0.74 ± 0.01 | **dA:rU** | > 5 | 0.96 ± 0.01 | **dA:rU** | 0.26 ± 0.04 | 0.84 ± 0.03 |
| **dG:rA** | ND | ND | **dG:rA** | slope = 0.006 ± 0.001 | | **dG:rA** | ND | ND |
| **dG:rG** | 0.04 ± 0.03 | 0.2 ± 0.1 | **dG:rG** | ND | ND | **dG:rG** | ND | ND |
| **dG:rC** | 2.4 ± 0.6 | 0.80 ± 0.02 | **dG:rC** | > 5 | 0.97 ± 0.01 | **dG:rC** | 0.42 ± 0.04 | 0.89 ± 0.02 |
| **dG:rU** | 0.52 ± 0.04 | 0.15 ± 0.01 | **dG:rU** | 0.11 ± 0.01 | 1.00 ± 0.02 | **dG:rU** | 0.21 ± 0.06 | 0.15 ± 0.01 |
| **dC:rA** | 0.6 ± 0.4 | 0.015 ± 0.002 | **dC:rA** | 0.22 ± 0.04 | 0.54 ± 0.03 | **dC:rA** | slope = 0.004 ± 0.001 | |
| **dC:rG** | > 5 | 0.80 ± 0.01 | **dC:rG** | > 5 | 0.95 ± 0.01 | **dC:rG** | 0.32 ± 0.02 | 0.87 ± 0.01 |
| **dC:rC** | 0.03 ± 0.01 | 1.0 ± 0.3 | **dC:rC** | slope = 0.028 ± 0.001 | | **dC:rC** | slope = 0.011 ± 0.001 | |
| **dC:rU** | 0.6 ± 0.2 | 0.08 ± 0.01 | **dC:rU** | 0.45 ± 0.07 | 0.87 ± 0.03 | **dC:rU** | 0.06 ± 0.02 | 0.21 ± 0.05 |
| **dT:rA** | 3.3 ± 0.5 | 0.69 ± 0.01 | **dT:rA** | > 5 | 0.93 ± 0.01 | **dT:rA** | 0.8 ± 0.1 | 0.76 ± 0.03 |
| **dT:rG** | 0.06 ± 0.02 | 0.4 ± 0.1 | **dT:rG** | 1.1 ± 0.2 | 0.61 ± 0.02 | **dT:rG** | slope = 0.012 ± 0.001 | |
| **dT:rC** | 0.04 ± 0.01 | 1.1 ± 0.2 | **dT:rC** | 0.12 ± 0.01 | 1.06 ± 0.02 | **dT:rC** | slope = 0.012 ± 0.001 | |
| **dT:rU** | 0.45 ± 0.07 | 0.19 ± 0.01 | **dT:rU** | 0.40 ± 0.04 | 0.89 ± 0.02 | **dT:rU** | 0.17 ± 0.04 | 0.08 ± 0.01 |

OT-2 HiKER Time Course Script
Written in Python

```
from opentrons import protocol_api
from opentrons.types import Point
import math

metadata = {
   'protocolName': '12.08.2023  P1-P4 Time Courses',
   'author': 'Zach <>',
   'apiLevel': '2.14'
}
#Performs multiple RNAP misincorporation experiments (for single plate, use RNAP Single protocol) in
sequence
#Place NTPs in first x rows, place enzyme in last x rows
#Will stop after 2 plates to prompt trash removal

#Number of Experiments (max 4, min 2)
exps = 4
#Pick 12 time points (in minutes)
timepoints = [0, 0.01, 0.5, 1, 2, 5, 7.5, 10, 12.5, 15, 17.5, 20]
def run(ctx):

   #SET-UP

   #load pipettes + tips
   #300 ul pipette and tips [RIGHT MOUNT, SLOT 11]
   tips300 = [ctx.load_labware("opentrons_96_tiprack_300ul", 11, "Tips300")]
   m300 =ctx.load_instrument('p300_multi_gen2', 'left', tip_racks = tips300)
   #20 ul pipette and tips [LEFT MOUNT, SLOTS 7, 4, 10, 2]
   tips20 = [ctx.load_labware("opentrons_96_tiprack_20ul", 7, "Tips20A"),
ctx.load_labware("opentrons_96_tiprack_20ul", 4, "Tips20B")]
   if exps >= 3:
      tips20 = tips20 + [ctx.load_labware("opentrons_96_tiprack_20ul", 10, "Tips20C")]
      if exps == 4:
         tips20 = tips20 + [ctx.load_labware("opentrons_96_tiprack_20ul", 2, "Tips20D")]
   m20 = ctx.load_instrument('p20_multi_gen2', 'right', tip_racks = tips20)

   #temp module [SLOT 1, 25C]
   temp_mod = ctx.load_module("temperature module gen2", 1) #[DO NOT CHANGE SLOT, ROBOT WILL
COLLIDE DURING CALIBRATION]
   react = temp_mod.load_labware('vwr_96_aluminumblock_100ul', label = "Reaction")  #Change to
'vwr_96_aluminumblock_100ul' to use vwr 100 uL pcr plate on temp mod
   temp_mod.set_temperature(celsius=25)

   #load plates [SLOTS 5, 6, 8, 9]
   finalA = ctx.load_labware('nestedcemult1_96_wellplate_300ul', 5, "FinalPlateA") #Change to
'nestedcemult1_96_wellplate_300ul' to use nested CE plate on biorad 96 well plate
   finalB = ctx.load_labware('nestedcemult1_96_wellplate_300ul', 6, "FinalPlateB") # ''
   FinalPlates = [finalA, finalB]
   if exps >= 3:
      finalC = [ctx.load_labware('nestedcemult1_96_wellplate_300ul', 8, "FinalPlateC")] # ''
```

```
    FinalPlates = FinalPlates + finalC
    if exps >= 4:
        finalD = [ctx.load_labware('nestedcemult1_96_wellplate_300ul', 9, "FinalPlateD")] # ''
        FinalPlates = FinalPlates + finalD

#water set up [SLOT 3]
reservoir = ctx.load_labware('nest_12_reservoir_15ml', 3,"Water")
water = [reservoir.wells()[n] for n in range(exps)] #Fill corresponding number of wells from L to R with water

#liquid set up
NTP = [react.rows()[0][n] for n in range(exps)] #Put NTP in first x rows
Enzyme = [react.rows()[0][n+8] for n in range(exps)] #Put enzymes in x rows, starting from row 9
FinalWells = []
for n in range(exps):
    FinalWells = FinalWells + [FinalPlates[n].rows()[0]] #Create list of wells in CE plates

#DEFINE FUNCTIONS
#time point function
#Old method that doesn't work with trash shoot
#def taketp(rxn,tp):
 #   m20.transfer(5, Enzyme[rxn], FinalWells[rxn][tp], mix_after = (2,10))


center_trash_location = ctx.fixed_trash['A1']
adjusted_trash = center_trash_location.top(z=55).move(Point(x=65, y=5))


#time point function
#Method of collecting time points that allows using the ramp

def taketp(rxn,tp,trash_location=adjusted_trash):
    m20.pick_up_tip()
    m20.aspirate(5,Enzyme[rxn])
    m20.dispense(5,FinalWells[rxn][tp])

    #One Mixing Step
    #m20.aspirate(10, FinalWells[rxn][tp], rate = 1)
    #m20.dispense(10, FinalWells[rxn][tp], rate = 1)
    #m20.drop_tip(trash_location)

    #unmute for Multiple Mixing Steps
    for rep in range(2):
        m20.aspirate(10, FinalWells[rxn][tp], rate = 1)
        m20.dispense(10, FinalWells[rxn][tp], rate = 1)
    m20.drop_tip(trash_location)




#master function to run assay
def runplate(tps, plate,trash_location=adjusted_trash):
```

```
        #time point 0
        #m20.transfer(2, Enzyme[plate], FinalWells[plate][0], mix_after = (2,7))
        m20.pick_up_tip()
        m20.aspirate(2,Enzyme[plate])
        m20.dispense(2,FinalWells[plate][0])
        for rep in range(1):
            m20.aspirate(7, FinalWells[plate][0], rate = 1)
            m20.dispense(7, FinalWells[plate][0], rate = 1)
            #m20.drop_tip(tip_racks["A12"].top(z=10))
        m20.drop_tip(trash_location)

        #begin rxn
        m300.transfer(50, NTP[plate], Enzyme[plate], mix_after = (2, 75))


        #further time points
        for n in range(len(tps)-1):
            ctx.delay(minutes = (tps[n+1]-tps[n]))
            taketp(plate,n+1)

    #BEGIN PROTOCOL
    for n in range(exps):
        runplate(timepoints,n,adjusted_trash)
        #if n == 1:
        #   ctx.pause('Empty Trash')

    #dilute with water
    for m in range(exps):
        m300.distribute(100, water[m], [FinalWells[m][n].top() for n in range(len(FinalWells[m]))], new_tip = 'once',
disposal_volume = 5)

    #deactivate heat module
    temp_mod.deactivate()
```

#### HiKER Exact Time Point Run Log Processing Script
#### Written in Python

```
import json
from datetime import datetime
import itertools
#Put in name of run log file
with open("11.22.2023 Exp1 RunLog.json", "r") as log:
    run = json.load(log)
Commands = run["commands"]["data"]
point_sample = 0
point_plate = 0

#Enter volume removed for each  time points (must be different than amount used to initiate reaction)
tpvol = 10

#Enter number of 96 well plates
Plates = 4
```

```
#Function which converts time readout to min
def get_min(time_str):
    # split in hh, mm, ss
    hh, mm, ss = time_str.split(':')
    return int(hh)*60 + int(mm)  + float(ss)/60


#Find Time Point 0 in run log file
PlateStarts = {}
for n in range(len(Commands)):
    if Commands[n]["commandType"] == 'dispense':
     if Commands[n]["params"]['volume'] == 50:
        PlateStarts[point_plate] = Commands[n]["completedAt"]
        point_plate = point_plate + 1


#Find Time Points 1+ in run log file
Times = {}
for n in range(len(Commands)):
    if Commands[n]["commandType"] == 'dispense':
        if Commands[n]["params"]['volume'] == tpvol:
            Times[point_sample] = Commands[n]["completedAt"]
            point_sample = point_sample + 1


#Convert Times to datetime
for n in Times:
    Times[n] = Times[n][:len(Times[n])-6]
    Times[n] = datetime.strptime(Times[n], '%Y-%m-%dT%H:%M:%S.%f')

#print(Times)
#print('\n\n')

#Index times for each plate to start
for n in PlateStarts:
    PlateStarts[n] = PlateStarts[n][:len(PlateStarts[n])-6]
    PlateStarts[n] = datetime.strptime(PlateStarts[n], '%Y-%m-%dT%H:%M:%S.%f')

#print(PlateStarts)


#Find True time points from when plate started
TimesBetween = []
for i in range(Plates):
    for n in range(11*i,11*i+11):
        TimesBetween = TimesBetween + [Times[n]-PlateStarts[i]]
```

```
#Print Difference between timepoints
TimeStrs = {}
for n in range(len(TimesBetween)):
   TimeStrs[n] = [str(TimesBetween[n])]

#print(PlateStarts)
#print(TimeStrs)

#Make CSV output file. Result is one column with 11 reaction time points for each plate. The first is exculded as it's zero.
#The reaction time stars back to zero when a new plate is started.
with open('Run Times.csv', 'w') as f:
   for key in TimeStrs.keys():
      f.write("%s, %s\n" % (key, get_min(''.join(TimeStrs[key]))))
```

HiKER Exact Time Point Run Log Processing Script
Written as Jupyter Notebook

```
#Importing Needed Tools

import numpy as np
from scipy.optimize import curve_fit
import matplotlib.pyplot as plt

#Import Data from Excel and Create Data Frame

import pandas as pd
data_df = pd.read_excel(r"C:\Users\zcarter\HiKER High-Throughput Calculation of kobs Script\11.22.2023 P4 Data_Analysis.xlsx",sheet_name ="Fin_Mat")


#Convert Data Frame to Arrary

exp_mat = pd.DataFrame(data_df).to_numpy()


#Label Individual Time Courses
```

```
t1 = exp_mat[:,0]
y1 = exp_mat[:,1]
t2 = exp_mat[:,2]
y2 = exp_mat[:,3]
t3 = exp_mat[:,4]
y3 = exp_mat[:,5]
t4 = exp_mat[:,6]
y4 = exp_mat[:,7]
t5 = exp_mat[:,8]
y5 = exp_mat[:,9]
t6 = exp_mat[:,10]
y6 = exp_mat[:,11]
t7 = exp_mat[:,12]
y7 = exp_mat[:,13]
t8 = exp_mat[:,14]
y8 = exp_mat[:,15]



# Define function you want to fit your data with

def func(x, k, A):
   return A *(1 - np.exp(-k * x))

#Fit your data

params1, pcov1 = curve_fit(func, t1, y1)
params2, pcov2 = curve_fit(func, t2, y2)
params3, pcov3 = curve_fit(func, t3, y3)
params4, pcov4 = curve_fit(func, t4, y4)
params5, pcov5 = curve_fit(func, t5, y5)
params6, pcov6 = curve_fit(func, t6, y6)
params7, pcov7 = curve_fit(func, t7, y7)
params8, pcov8 = curve_fit(func, t8, y8)



# Report Best Fit Parameters (kobs, Amp) and Generate Best Fit Lines With Experimental Data

print(params1)
print(params2)
print(params3)
print(params4)
print(params5)
print(params6)
print(params7)
print(params8)




plt.scatter(t1, y1)
plt.scatter(t2, y2)
```

```
plt.scatter(t3, y3)
plt.scatter(t4, y4)
plt.scatter(t5, y5)
plt.scatter(t6, y6)
plt.scatter(t7, y7)
plt.scatter(t8, y8)



xsim = np.linspace(0, 28, 150)

plt.plot(xsim, func(xsim, *params1))
plt.plot(xsim, func(xsim, *params2))
plt.plot(xsim, func(xsim, *params3))
plt.plot(xsim, func(xsim, *params4))
plt.plot(xsim, func(xsim, *params5))
plt.plot(xsim, func(xsim, *params6))
plt.plot(xsim, func(xsim, *params7))
plt.plot(xsim, func(xsim, *params8))


#Organize simulations into master array
sim_master_array=np.array([xsim, func(xsim, *params1),xsim, func(xsim, *params2),xsim, func(xsim,
*params3),xsim, func(xsim, *params4),xsim, func(xsim, *params5),xsim, func(xsim, *params6),xsim,
func(xsim, *params7),xsim, func(xsim, *params8)])
sim_master_array=np.transpose(sim_master_array)

#Convert master simulations array into a data frame
Sims_Matrix_DF = pd.DataFrame(sim_master_array,columns=['t1', 'y1', 't2', 'y2', 't3', 'y3', 't4', 'y4', 't5', 'y5', 't6',
'y6', 't7', 'y7', 't8', 'y8'])


#Export Fit Parameters, Experimental Data, and Simulated Data to Excel Sheet

#Orgnize fit parameters into one master array
Params_Matrix_Arr = np.array ([params1,params2,params3, params4, params5, params6, params7,
params8])

#Convert master fit parameters array into a data frame
Params_Matrix_DF = pd.DataFrame(Params_Matrix_Arr, columns=['kobs','Amp'])



with pd.ExcelWriter('Data_Analysis.xlsx') as writer:
    Params_Matrix_DF.to_excel(writer, sheet_name='Fit_Parameters')
    data_df.to_excel(writer, sheet_name='Experimental_Data')
    Sims_Matrix_DF.to_excel(writer, sheet_name='Simulation_Data')
```
